# Effect of Localization on the Stability of Mutualistic Ecological Networks

**DOI:** 10.1101/023275

**Authors:** S. Suweis, J. Grilli, Jayanth R. Banavar, S. Allesina, A. Maritan

**Affiliations:** Department of Physics and Astronomy, University of Padua, Consorzio Nazionale Interuniversitario per le Scienze Fisiche della Materia and Istituto Nazionale di Fisica Nucleare, 35131 Padova, Italy.; Department of Ecology and Evolution, University of Chicago, 1101 East 57th Street, Chicago, Illinois 60637, USA.; Department of Physics, University of Maryland, College Park, Maryland 20742, USA.

## Abstract

The relationships between the core-periphery architecture of the species interaction network and the mechanisms ensuring the stability in mutualistic ecological communities are still unclear. In particular, most studies have focused their attention on asymptotic resilience or persistence, neglecting how perturbations propagate through the system. Here we develop a theoretical framework to evaluate the relationship between architecture of the interaction networks and the impact of perturbations by studying localization, a measure describing the ability of the perturbation to propagate through the network. We show that mutualistic ecological communities are localized, and localization reduces perturbation propagation and attenuates its impact on species abundance. Localization depends on the topology of the interaction networks, and it positively correlates with the variance of the weighted degree distribution, a signature of the network topological hetereogenity. Our results provide a different perspective on the interplay between the architecture of interaction networks in mutualistic communities and their stability.

## Introduction

Ecological networks may be viewed as a set of species (nodes) connected by interspecific interactions (competition, predation, parasitism, and mutualism), represented by the links. Even though interaction strengths are largely unknown, the architecture of the ecological interaction networks has been thoroughly investigated, showing its important role in shaping and regulating community dynamics and in structuring diversity patterns ^1–8^. Several studies recognized the strong impact of the non-random structures of empirical interaction networks on both the resilience (time to return to the steady state after a small perturbation) and the persistence (number of coexisting species at equilibrium) of ecological communities ^9–14^, and much theoretical effort has been made to understand the relationship between stability and complexity in ecological communities, one of the most debated issues in ecology ^15–18^. In mutualistic networks, where species beneficially interact with each other, a core-periphery structure has been observed ubiquitously ^19^. The network core refers to a central and densely connected set of nodes, while the periphery denotes a sparsely connected non-central set of nodes, which are linked to the core. It has been posited that the architecture of mutualistic networks minimizes competition and increases biodiversity ^7^, community stability (resilience) and persistence ^20^, but other studies have demonstrated that structured mutualistic ecological networks may be less stable than their random counterparts ^14,21^. It has also been shown that community stability decreases as community size increases, and that this result holds even for more realistic ecological interactions with a mixing of interaction types (“hybrid communities”) ^22^. Most of the aforementioned studies focused either on the resilience of the system - measured by the maximum real part of the eigenvalues of the community matrix ^14,15,21^ or on the number of species that persist when starting from non-stationary conditions ^7,8^. However, both approaches have important limitations. Indeed, the maximum real part of the community matrix eigenvalues only describes the rate of recovery from perturbations in the long time limit, providing no information on the transient response. Perturbations can grow significantly before decaying, possibly impacting species’ fate (see Figure 1A). A system at its stable stationary state that experiences such initial amplifications of the perturbations is called reactive ^23,24^. On the other hand, persistence (measured as the fraction of initial species with positive stationary population density ^16^) is strongly sensitive to initial conditions, the system’s distance from stationarity and the choice of model and parameters ^8,25,26^. To garner a better understanding of the effect of perturbations on ecological communities, one should also study how the components of the leading eigenvectors (i.e., the right and left eigenvectors associated with the eigenvalue having the largest real part) are distributed, i.e., study the localization of the system. In condensed matter physics, localization, also known as Anderson localization ^27^, is the absence of diffusion of waves in a disordered medium, and it describes the ability of waves to propagate through the system. Other approaches (e.g. Markov chain models ^28^, or the inverse community matrix ^29^) can be used to study how disturbances propagate in species interaction networks and what their effects are on other species (i.e. how many other species do they affect and what is the magnitude of this effect). However, it has been shown that small variations in the interaction strengths may lead to very different model predictions ^30,31^. Our theoretical framework may be considered as a complementary methodology to gain information on the general relation patterns between the interaction network architecture and the ability of perturbations to propagate within the system. Our goal in this work is to determine the degree of localization of eigenvectors in mutualistic ecological networks as a function of the network size, structure and interaction strengths, and to study the impact of localization on the perturbation amplitude and spreading within the system. Here we show that localization may be a useful mechanism that impacts on the stability of ecological networks. In fact, localization attenuates (asymptotically) and reduces perturbation propagation through the network. We find that mutualistic ecological networks are indeed localized and localization patterns are correlated with some network topological properties; in particular, heterogeneity in the weighted species degrees promotes localization in the network. Furthermore, the observed localization increases with the size of the ecological communities, highlighting a trade-off between the asymptotic resilience of the system and the attenuation of perturbations.

**Figure 1:**
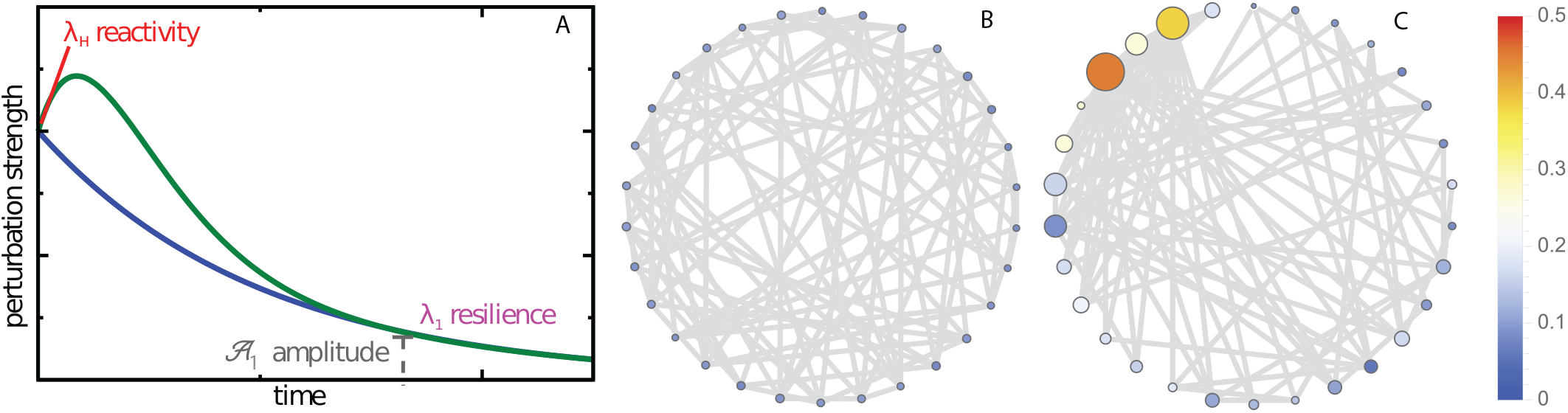
Propagation of the perturbation through the network. A) Trajectory of a perturbation through time. Reactivity (*λ*_*H*_) measures whether perturbations grow before decaying; Asymptotic resilience *λ*_1_ indicates whether perturbations eventually decay; and the asymptotic perturbation amplitude *𝒜*_1_ describes the intensity of the perturbation for large time. The principal right eigenvector determines which species will be affected most by the perturbation after its propagation, while the left principal eigenvector controls which species are the most sensitive to the initial perturbation (i.e. before its propagation - see Methods). The weighted degree heterogeneity affects the localization pattern in the network: B) is a regular graph where each node is connected to 6 other nodes, while C) is a power-law scale-free graph ^2^ of the same size and with similar connectance. In both cases, edge weights are randomly extracted from a Gamma distribution. The size and the color of the nodes indicate the absolute values of the corresponding component of the leading right eigenvector. In B, all species are equally perturbed. In contrast, in C, only few species are affected.

## Results

### Theoretical Framework

The mutualistic interactions of an ecological community can be encoded in a bipartite binary graph represented by its adjacency matrix *B* containing *S* nodes (species) that are partitioned into two disjoint sets, one containing the animals (insect pollinators), the other the plants. Each of the *L*(undirected) edges connects two nodes, one in the set of animals (of size *A*) and the other in the set of plants (of size *P*), i.e., *B*_*kl*_= 1 if insect *k* and plant *l* interact. *S*= *A* + *P* is the total number of species in the community. We analyze 59 bipartite binary networks available from the interaction web database ^9^, and we construct the *S × S* community matrix Φ describing the linearized system dynamics, by assigning to each animal-plant interaction a positive “weight” (see Methods). Let **x**(*t*) be the *S*-component vector describing the abundance of the *S* populations at time *t*. The propagation of a given small perturbation ***ξ***=(*ξ*_1_, *ξ*_2_, *…, ξ_S_*) acting on the system at stationarity will lead to small departures, *δ***x**(*t*), from the stationary state **x**\* and can be studied by the linearized system of coupled differential equations 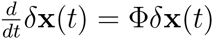, where *δ***x**(*t*) = **x**(*t*)*-***x***\** (with *δ***x**(0) =***ξ***, which in turn can be studied in terms of the eigenvectors and eigenvalues of Φ, known as community matrix (see Methods). In particular, the asymptotic behavior of the perturbed systems can be analyzed in terms of the largest eigenvalue *λ*_1_ and corresponding left and right eigenvectors **u**_1_ and **v**_1_, i.e., 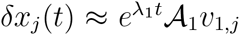 for large *t*, where *𝒜*_1_ = (**u**_1_ *·****ξ***)/(**v**_1_ *·***u**_1_) is the amplitude associated with the asymptotic propagation of the perturbation through the ecological network (Figure 1 and Methods).

Clearly, the relaxation of a system to its equilibrium state after a perturbation is not uniquely controlled by the leading eigenvalue of the community matrix. All the eigenvalues contain information on the time-scales involved in the relaxation, while the corresponding eigenvectors determine how the perturbation spreads and relaxes in different species. The leading right eigenvector, in particular, sets the relative vulnerability of species and how they are affected by perturbations in the long run. If this eigenvector is localized, i.e., if only few components/species have non negligible values, then a perturbation after its propagation involves only few species. On the other hand, the left leading eigenvector indicates which species are most hit by the perturbation before its propagation. It also plays an important role in modulating the amplitude of the perturbation (that is proportional to ***ξ****·***u**_1_ see Methods). Consider for example a 4 × 4 community matrix for which, 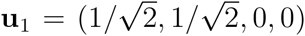 and 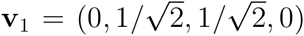. If ***ξ***=(0,0,1,1), then **u**_1_ *ξ*= 0 and the perturbation decay time will be very fast (controlled by *λ*_2_, rather than *λ*_1_ see Eq.(1) in the Methods section). On the other hand, if ***ξ***= (1,1,0,0), then the system asymptotic recovery time will be longer (proportional to 1*/λ*_1_), but only species 2 and 3 will be affected at these time scales. As a general trend, we will show that localization mainly depends on the heterogeneity of the network weighted degrees (or strengths **s** = (*s*_1_, *s*_2_, *…, s_S_*)): in the case of high variability in these strengths, the system display localization (see Figure 1B-C). The behavior immediately after the perturbation (i.e., in the limit *t* → 0^+^) can be analyzed by studying its reactivity ^23^, defined as the maximum amplification rate over all initial perturbations, and immediately after the perturbation. It can be shown ^23^ that the reactivity *λ*_*H*_can be computed as the maximum eigenvalue of *H*= (Φ + Φ^*T*^)/2, the symmetric part of Φ. If *λ*_1_ *<*0 and *λ*_*H*_ >0 then the equilibrium point is stable but reactive. Because *λ*_*H*_ ≥ *λ*_1_ ^23^, the reactivity can also be used to develop an early warning signal for systems approaching a non stable stationary state ^24^. If the eigenvector **w**_*H*_corresponding to *λ*_*H*_, is also localized, then it means that the perturbation magnitude on these localized species will tend to grow (see Figure 1A), i.e. in the short time, these species will be the most affected by the perturbation.

### Localization Patterns

We compare localization patterns of 59 empirical mutualistic networks, and two corresponding random null models. In the first null model, we randomize the interactions while keeping the networks connected. In the second null model, we randomize the interactions, but we also constrain the network degrees sequence *{k*_1_, *k*_2_, *…, k_S_}* to be as in the corresponding empirical networks (see Methods). To measure localization, we use the inverse participation ratio (*IPR*) ^27^, the classical way to quantify how many relevant components are observed in the leading eigenvectors (see Methods). The degree of localization increases as *IPR* increases. If *IPR* is one, then only one component of the eigenvector is non-zero. We quantify the presence of localization by computing the *rIPR* defined as the ratio between the *IPR* of each real empirical network and the *IPR* of the corresponding random null model.

As Figure 2A–B shows, most of the empirical networks are significantly more localized in both the right and left leading eigenvectors with respect to null model 1, while they have the same level of localization of null model 2 (Figure 2D–E,I and Table 1). These results suggest that it is the core-periphery network structure of empirical systems (a manifestation of heterogeneous degree distributions) that is responsible for their higher localization: once we constrain the degree distributions to be fixed (that in the case of mutualistic networks are most likely approximate truncated power laws ^2^), then null model 2 generates localization patterns very similar to those observed in empirical mutualistic networks (see Table 1). Nodes strength *s*_*i*_ (or weighted degrees) also play a crucial role. In fact, an adjacency network with core-periphery structure, but having ‘anti-nested’ distributed weights ^13^, will not be localized because, contrary to its degree distributions, the weighted degree distribution will be homogeneous (see also Supplementary Information, section 5). The localization of **w**_*H*_ displays the same patterns (Figure 2C,F), and we found that species that are the most affected by the perturbation at short time scales are also those that absorb most of the perturbation asymptotically - indicating a limiting capability of the the perturbation to propagate through the network. In fact, the position of the localized components for **w**_*H*_ is most likely to be the same of those for **v**_1_ and **u**_1_ (see Figure 3 and Supplementary Information, section 7).

**Table 1:** Statistics of Localization Patterns. Statistics of localization patterns for three different ecological scenarios (described by δ - see Methods) given by the fraction of localized empirical networks with respect to generated null models (NM) for different parametrization (*rIPR* > 1 and *P*-value < 0:05) and corresponding asymptotic attenuation of the perturbation 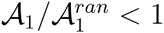 and *P*-value< 0:05). We found that most of the empirical networks are indeed localized with respect to NM1, but not with respect to NM2.

| $IPR/IPR - \text{ran} < 1$ | $\delta = 0.5$ | $\delta = 0$ | $\delta = -0.5$ | NM # |
| --- | --- | --- | --- | --- |
| $\mathbf{u}_1$ | $\approx 86\%$ | $\approx 76\%$ | $\approx 39\%$ | 1 |
| $\mathbf{v}_1$ | $\approx 61\%$ | $\approx 64\%$ | $\approx 42\%$ | 1 |
| $\mathbf{w}_H$ | $\approx 83\%$ | $\approx 72\%$ | $\approx 50\%$ | 1 |
| $\mathbf{u}_1$ | $\approx 20\%$ | $\approx 20\%$ | $\approx 19\%$ | 2 |
| $\mathbf{v}_1$ | $\approx 27\%$ | $\approx 27\%$ | $\approx 11\%$ | 2 |
| $\mathbf{w}_H$ | $\approx 24\%$ | $\approx 20\%$ | $\approx 19\%$ | 2 |
| $\mathcal{A}_1/\mathcal{A}_1^{ran} < 1$ | $\delta = 0.5$ | $\delta = 0$ | $\delta = -0.5$ | NM # |
| | $\approx 81\%$ | $\approx 81\%$ | $\approx 70\%$ | 1 |
| | $\approx 17\%$ | $\approx 15\%$ | $\approx 14\%$ | 2 |

**Figure 2:**
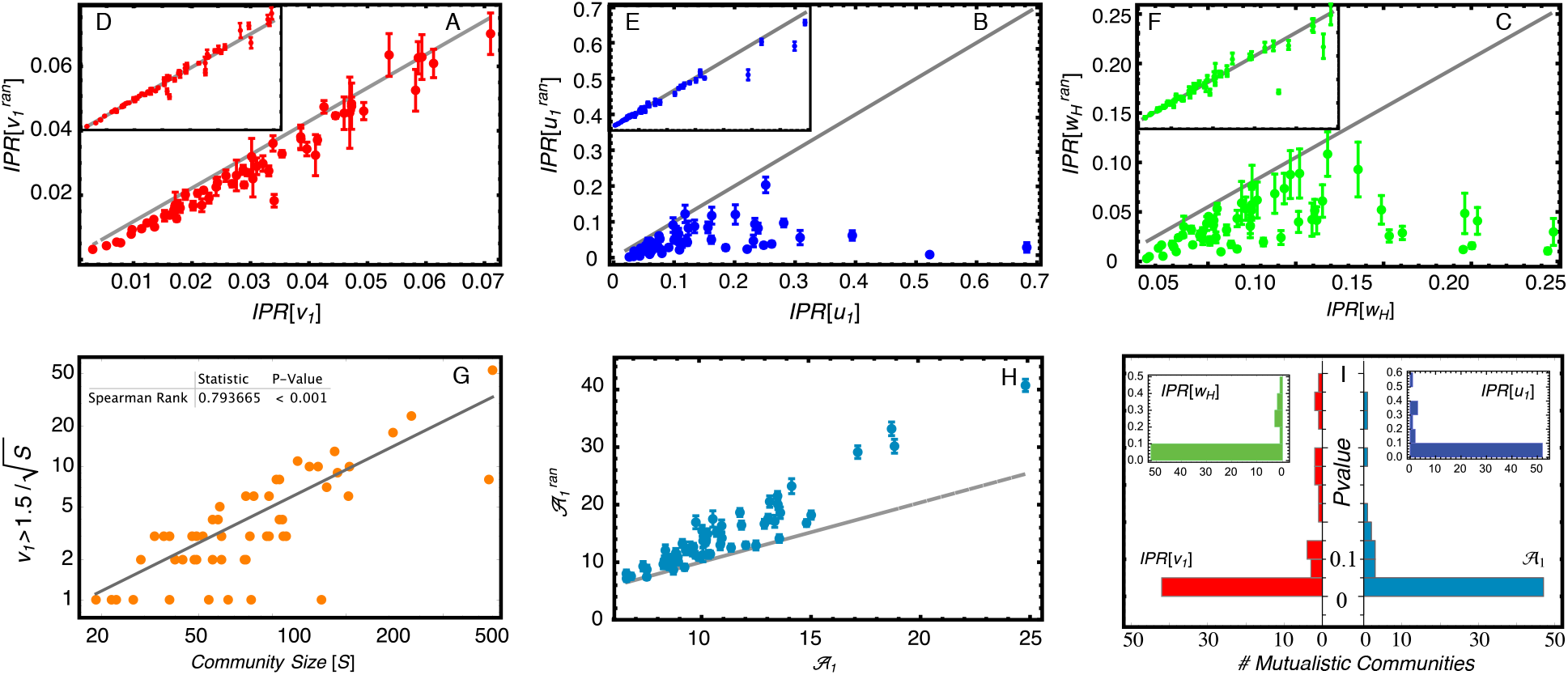
Localization Patterns and Effect on the Asymptotic Amplitude. A-C) Localization (*IPR* see Methods) of the leading eigenvectors for null model 1 versus empirical mutualistic networks (right eigenvector **v**_1_ in red, left eigenvector **u**_1_ in blue and reactive eigenvector in green). Points represent average value of 1000 randomizations, bars indicate the standard deviation. 1:1 line represents the value of the empirical mutualistic networks. D-F) Same for null model 2 (see Methods). G) Number of (localized) components as a function of the community size: a significant correlation is observed (Spearman Rank Test = 0.715). H) Effect of the localization on the asymptotic amplitude *𝒜*_1_ for the simulated perturbation *ξ*_*all*_ on empirical mutualistic networks with respect to null model 1 *𝒜* - ran. In *≈* 85% of the cases, the perturbation amplitude is significantly attenuated (*P – value ≤* 0.05 see also Table 1). I) *P - values* of the observed values of localization and asymptotic amplitudes in empirical networks with respect to null model 1. Parameters here are *δ* = 0.5 and *γ*_0_ = 1 (see Methods). For robustness of the results with respect to the parametrization see Supplementary Information, section 4.

**Figure 3:**
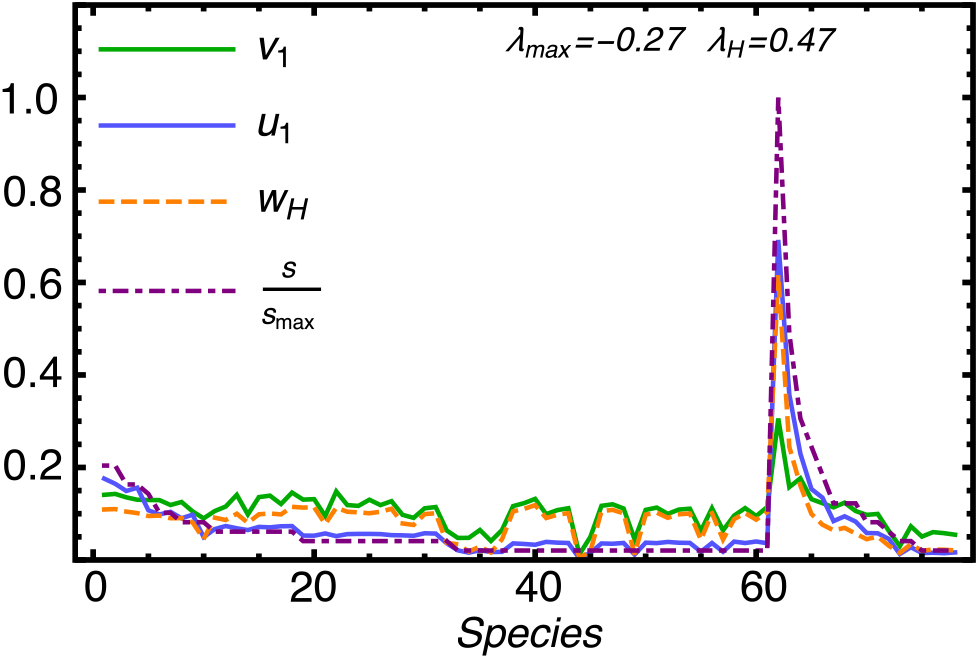
Localization in Norfolk mutualistic insect-grasslands community. Example of localization in a real mutualistic community ^35^ with *S* = 78 species. The 61 insects and the 17 plants species are sorted according to degree. The flower species *Leucanthemum vulgare* is the most localized in each of the three eigenvectors, **v**_1_, **u**_1_ and **w**_*H*_ and corresponds to the species with the highest species degree and strength. The parameters are *δ* = 0.5 and *γ*_0_ = 1, yielding a stable but reactive equilibrium (*λ*_1_ *<* 0, *λ*_*H*_ > 0).

### Relation between Localization and Network Topological Properties

Localization patterns in empirical mutualistic communities depend on both the size and the connectance of the species interaction network (Figures 2G and 4). While the leading eigenvalue *λ*_1_—the one with the largest real part which controls the relaxation time—increases for increasing community size ^15^ (assuming that *γ*_0_ does not scale ^22^ with *S*), we observe (Figure 4A) an interesting strong positive correlation between community diversity (network size) and localization (rIPR). We also note that in empirical pollination communities, networks size is negatively correlated with network connectance ^14^. Network connectance, in turn, is negatively correlated with localization (Figure 4B): the higher the connectance, the higher is the ability of perturbations to propagate through the network (a general property observed also in financial networks ^32^, and socio-environmental interdependent systems ^33,34^), and thus the lower the level of localization. The strong positive correlation between network size and localization leads to a trade-off between localization and asymptotic resilience (in terms of *λ*_1_ see Table 2). This result may shed light on the celebrated complexity-diversity paradox ^15,17^: the less an ecological community is resilient, the more it is localized, and asymptotic perturbation is attenuated (see Figure 5). We finally note that localization is positively correlated with the variance of the weighted degree distribution, see Figure 4C). This correlation reflects the fact the localization is a manifestation of the heterogeneity of the network topology. In the Supplementary Tables S1-S3, we also calculate the correlations between localization (*rIPR*) and the topological properties of the networks under different parametrizations and with respect to null model 2. We found that in this latter case the correlations are not significant. We can thus conclude that it is indeed the heterogeneity in the weighted degree distribution, which is the key structural aspect of ecological networks that is related to localization.

**Table 2:** Correlations between network topological and spectral properties. Correlations *ρ*(*x*; *y*) measured using Spearman Rank Test (parametrization *δ* = 0.5) using Holling Type I model with *a*=40 and *b*=0.05 - see Methods). rIPR refers to null model 1.

| $\rho(x, y)$ | $x$ | $y$ | $P - value$ |
| --- | --- | --- | --- |
| $\approx 0.760$ | $S$ | $rIPR[\mathbf{v}_1]$ | $< 10^{-4}$ |
| $\approx 0.754$ | $S$ | $rIPR[\mathbf{u}_1]$ | $< 10^{-4}$ |
| $\approx 0.800$ | $S$ | $rIPR[\mathbf{w}_H]$ | $< 10^{-4}$ |
| $\approx -0.662$ | $C$ | $rIPR[\mathbf{v}_1]$ | $< 10^{-4}$ |
| $\approx -0.754$ | $C$ | $rIPR[\mathbf{u}_1]$ | $< 10^{-4}$ |
| $\approx -0.749$ | $C$ | $rIPR[\mathbf{w}_H]$ | $< 10^{-4}$ |
| $\approx -0.769$ | $S$ | $\mathcal{A}_1$ | $< 10^{-4}$ |
| $\approx 0.806$ | $C$ | $\mathcal{A}_1$ | $< 10^{-4}$ |
| $\approx -0.477$ | $\lambda_1$ | $\mathcal{A}_1$ | $< 10^{-4}$ |
| $\approx 0.578$ | $\lambda_1$ | $rIPR[\mathbf{v}_1]$ | $< 10^{-4}$ |
| $\approx 0.390$ | $\lambda_1$ | $rIPR[\mathbf{u}_1]$ | $< 10^{-4}$ |
| $\approx 0.468$ | $\lambda_1$ | $rIPR[\mathbf{w}_H]$ | $< 10^{-4}$ |
| $\approx 0.460$ | $\sigma_s^2$ | $rIPR[\mathbf{u}_1]$ | $< 10^{-4}$ |
| $\approx 0.535$ | $\sigma_s^2$ | $rIPR[\mathbf{v}_1]$ | $< 10^{-4}$ |
| $\approx 0.45$ | Degree [k] | $\mathbf{v}_1$ | $< 10^{-4}$ |
| $\approx 0.46$ | Strength [s] | $\mathbf{v}_1$ | $< 10^{-4}$ |
| $\approx 0.67$ | Degree [k] | $\mathbf{u}_1$ | $< 10^{-4}$ |
| $\approx 0.61$ | Strength [s] | $\mathbf{u}_1$ | $< 10^{-4}$ |
| $\approx 0.76$ | Page Rank | $\mathbf{v}_1$ | $< 10^{-4}$ |
| $\approx 0.94$ | Eig. Centrality | $\mathbf{v}_1$ | $< 10^{-4}$ |

**Figure 4:**
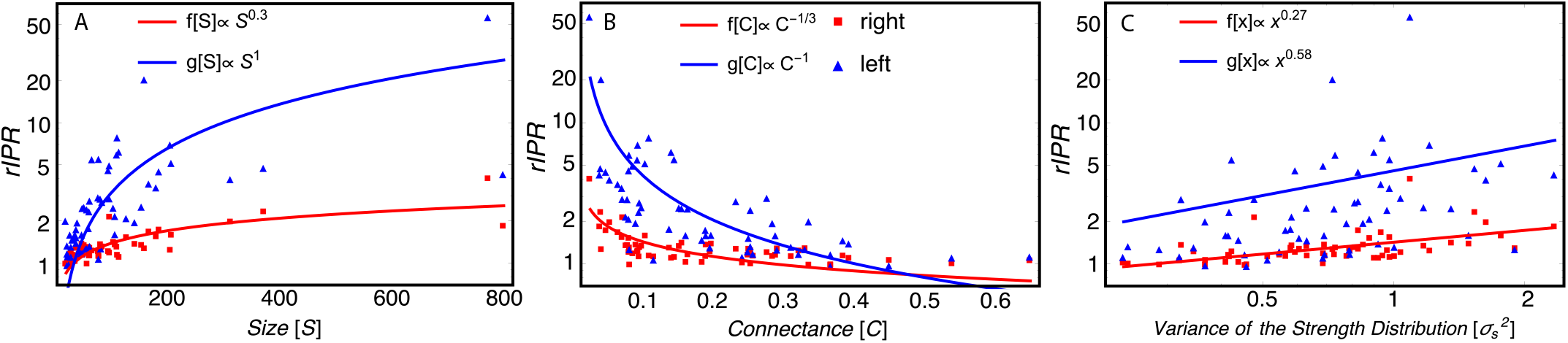
Relation between localization and network topological properties. Relation between localization (rIPR calculated through null model 1 see Methods) and A) networks size (S); B) network connectance measured as the fraction of observed and possible links (*C* = *L/*(*S*(*S -* 1)) with *L*=number of links); C) network heterogeneity (variance of the weighted degree distribution). The parametrization used here is *δ* = 0.5 and *γ*_0_ = 1. Results are reported for both right (in red) and left (in blue) leading eigenvectors (**v**_1_ and **u**_1_).

**Figure 5:**
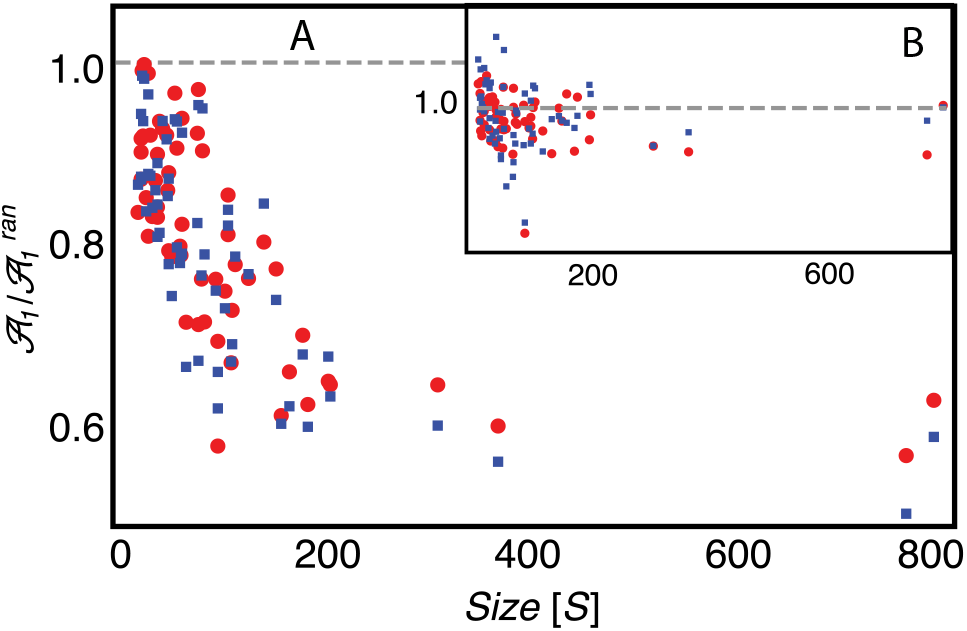
Relation between Size and Relative Asymptotic Amplitude. A) Relationship between the relative asymptotic amplitude (with respect to null model 1) and network size: attenuation increases for increasing community size (i.e., 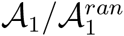 decreases with *S*). B) Asymptotic amplitude for perturbed empirical communities is compatible with that one generated by null model 2 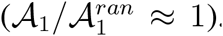. Indeed null model 2 generates networks with the same level of localization of the empirical pollinator networks (*rIPR ≈* 1 see Figure 2 D-F), and thus attenuation is not observed. Red points correspond to the parametrization *δ* = 0.5 and *γ*_0_ = 0.025, while blue squares represent the mean field case (*δ* = 0, *γ*_0_ = 1).

### Attenuation of Perturbation Propagation and Amplitude due to Localization

In order to make analytical progress and to better understand the impact of the localization on the amplitude of the perturbation spreading throughout the network, we analyze the mean field case (*δ* = 0), and we assume that the perturbation vector is a constant, i.e., ***ξ***= *ξ*_0_**1** (a *S* vector of unit components) and **v**_1_ *≈* **u**_1_. Under this assumption, we are able to prove that **𝒜**_1_ = |ξ_0_|(Σ_*j*_ |*v*_1,*j*_|)^2^ and thus *𝒜*_1_ is maximum when 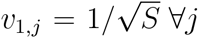 which corresponds to a state of minimum localization of Φ, while *A*_1_ is minimum when *v*_1,*j*_ = *δ*_*j,i*_ for some *i*, which corresponds to the fully localized case. Thus, the mean-field approximation with constant perturbation suggests that localization in the system reduces the amplitude of the principal mode of the perturbation wave. Indeed, our numerical simulations (Figures 2H and 5A and Supplementary Information, section 4.1) confirm that a localized structure leads to a decrease in the principal amplitude *𝒜*_1_ of the perturbation also beyond the mean-field case (i.e. *δ ≠* 0). Moreover, following the localization trend, the perturbation damping increases with the size of the system: the larger the ecological network, the stronger is the attenuation due to the system localization (Figure 2D). Also the reverse is true: if a network is not significantly more localized than its corresponding null model, then no attenuation is observed (see Figure 5B, Table 1 and Supplementary Information, section 4.2).

## Discussion

We have developed a comprehensive theoretical framework to evaluate the relationship between the species interaction network architecture and the impact of a given perturbation on ecological mutualistic networks. Localization has thus two beneficial effects on ecological network robustness: a) only a very low proportion of species in the community are significantly affected by a perturbation spreading throughout the network, and b) localization leads to an attenuation of the perturbation effects on the system. These results are robust with respect to variation of the parameters (see Supplementary Information, sections 2 and 4) and thus hold for very general parametrization of the interaction strengths (that in general are unknown see Supplementary Information, section 1.1). We thus have shown that the eigenvectors of the community matrix play a crucial role in determining the impact and the propagation of the perturbation through the system. We found that the positions of the localized components of the principal eigenvectors strongly correlates with nodes degree centrality, species strength *s*_*i*_, eigenvector centrality and page-rank centrality (Table 2 and Supplementary Information, section 7): the proposed framework thus allows one to identify those species which are affected the most by a given perturbation. Interestingly, these are species with many mutualistic interactions, and on average with higher population abundances with respect to specialist species ^14^. For example, in Figure 3 we show the eigenvectors components of the leading eigenvectors for the community and reactivity matrix associated with the insect-grasslands ecological community in Norfolk ^35^. For each of the 61 insects and the 17 plants, we can calculate the corresponding values of **v**_1_, **u**_1_ and **w**_*H*_. Species within each class (pollinators and flowers) are then sorted according to their degree (number of interacting partners they have). We note that the flower species *Leucanthemum vulgare*, the species with the highest species degree and a high density, is the most localized species in the community for both **v**_1_, **u**_1_, and **w**_*H*_, and it is the one that is likely to absorb most of a potential perturbation affecting the whole community.

A general emerging pattern observed for the mutualistic communities analyzed in this work is that, while these systems are less resilient for increasing biodiversity (May’s result ^15^), localization and the corresponding perturbation attenuation increase with increasing species diversity. In other words, these mutualistic systems experience a trade-off between resilience and localization: small communities are faster in recovering their stable state after a perturbation, but they are less localized and the perturbation will have an impact on most of the species. On the other hand, large communities are less resilient (i.e., need larger time to return to a stable state), but only few species will be affected by the perturbation, and amplitude of the perturbation will be attenuated while spreading through the network. The proposed theoretical analysis can be easily applied to networks with other interaction types, and it is illustrative of the potential of this new metric.

## Methods

### Data Parametrization

In order to be able to describe different ecological scenarios, and following recent models in the literature ^26,36^, we parameterize the weighted interaction matrix as 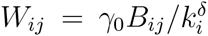 for *i ≠ j* and *W*_*ii*_ = *−d_i_* (see Supplementary Information, section 1), where *B* is the adjacency matrix of the species interaction matrix, indicating presence (*B*_*ij*_=1) or absence (*B*_*ij*_=0) of interactions among species, *k_i_* = ∑_*i*_*B_ij_* is the number of *mutualistic* partners of species *i* (species degree), *s_i_* = ∑_j_*W_ij_* is the species strength (or weighted degree) and *γ*_0_ is a parameter describing the basal mutualistic strength, while *δ* a trade-off parameter controlling the relation between mutualistic interaction strength and species degree. Following a Holling Type I population dynamics model, we then build the community matrix Φ as 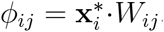, where 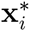 denotes the stationary population abundance of species *i*, and we model it as random variable drawn from a Gamma distribution so to have an average species population abundance 〈*x*\*〉 = 1 and of standard deviation *σ*_*x*_* = 0.158 (see Supplementary Information section 1). By varying the parameter *δ* we investigate: a) The architecture with constant interaction strength (“mean-field” case ^7,14^, *δ* = 0); b) The architecture with interaction strength-degree trade-off (*δ >* 0), e.g. specialist species interact stronger than the generalists one) ^37,38^. c) Architecture where generalist species interact stronger than specialist species (*δ <* 0). Using this parametrization, for a fixed *δ* the stability and reactivity of ecological communities can be controlled by the value of the basal mutualistic strength *γ*_0_, and the intra-specific competition *d*_*i*_.

### Null Models

We generate two different random null models (NM) for Φ (Φ – ran). NM1) We assign the *L* links in the adjacency matrix at random while keeping the network connected, and then parametrize it in the same way we do for empirical networks. NM2) We assign the *L* links at random, but constraining the degree sequence (*k*_1_, *k*_2_, *…, k_S_*) to be the same of the corresponding adjacency matrix *B* and then parametrize it in the same way we do for empirical networks. Our results are compared with 1000 realizations of each of the null models. For all other details we refer to the Supplementary Information, section 3.

### Perturbation Analysis

The effect of a given perturbation ***ξ***= (*ξ*_1_, *ξ*_2_, *…, ξ_S_*) acting on the system at time *t* = 0 will propagate in time obeying 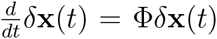 with initial condition **x**(0) =***ξ***. The solution of the latter equation can be written in terms of the eigenvectors and eigenvalues of Φ:

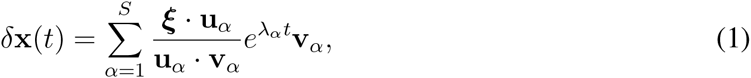

where **u**_*α*_, **v**_*α*_ and *λ*^(*α*)^ are respectively the left, the right eigenvectors and the corresponding eigenvalues of the linearization matrix Φ. We ordered the eigenvalues so that 0 *> λ*_1_ *>* Re [*λ*^(^2^)^]*>*… *>* Re [*λ*^(*n*)^](we note that in our case, as *ϕ*_*ij*_ ≥ 0, the Perron-Frobenius theorem holds and *λ*_1_ = Re [*λ*^(^1^)^]). For simplicity, we will denote by *𝒜*_*α*_ = (***ξ*** *·* **u**_*α*_)/(**u**_*α*_ · **v**_*α*_) the amplitude associated with the *α*-*th* mode of the perturbation.

### Localization and Effect on Stability

We measure the localization using the inverse participation ratio 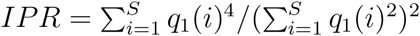, where **q**_1_ = **v**_1_, **u**_1_ or **w**_*H*_. In particular, we identify localization patterns by computing the *rIPR*, the ratio between the *IPR* of each real empirical network and the *IPR* of the corresponding random null model: *rIPR*_i_ = *IPR_i_*/〈*IPR_i_^ran^*〉. The average 〈·〉 is taken among different realizations of Φ - ran. If *rIPR* is significantly larger than one, then the system is localized. Otherwise we say that the system is not localized. We can also quantify the number of localized species by setting a threshold *θ* and count the fraction of species with a leading eigenvector component larger than that threshold, i.e *v*_1_(*i*) *> θ* or *u*_1_(*i*) *> θ*. We set 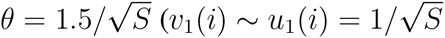 would correspond to the extended, non localized case). We quantify how the architecture of the ecological networks affects the impact of a simulated perturbation on the system by comparing the outputs *𝒜*_1_, *λ*_1_, *λ*_*H*_, **v**, **u**, **w**_*H*_ and *ρ* with respect to the corresponding random null models. We consider different type of perturbations. In Supplementary Information (section 6), we present results for *a*) A noise ***ξ***_*D*_ which is independent of species characteristics, i.e. ***ξ***_*D*_ drawn from a normal distribution *𝒩* (1, *ζ*) of mean 1 and variance *ζ*^2^; b) A noise ***ξ***_*E*_ that is species dependent, i.e., proportional to the degree of each species (*ξ*_*E*_(*i*) *∝ k_i_ξ_D_*(*i*)). In the main text we show results for a perturbation combining both types of noise, i.e., ***ξ***_all_ =***ξ***_*D*_+***ξ***_*E*_. A link to the Mathematica notebook with main functions needed to compute localizations and effect on stability of mutualistic ecological communities is here provided: https://github.com/suweis/Effect-of-Localization-on-the-Stability-of-Mutualistic-Ecological-Networks

## Acknowledgements

SS thanks the University of Padova, Physics and Astronomy Department Senior Grant 129/2013 Prot. 1634, JG is funded by the Human Frontier Science Program; AA is grateful to PRIN 2012, SA supported by NSF DEB #1148867.

## Competing Interests

The authors declare that they have no competing financial interests.

## Correspondence

Correspondence and requests for materials should be addressed to SS.

## Author contributions

S.S. J.G, S.A, J.R.B., and A.M. designed research; S.S., and J.H. performed research and numerical simulations; and S.S. J.G, S.A, J.R.B., and A.M. wrote the paper.

